# An endogenous opioid circuit determines state-dependent appetitive behavior

**DOI:** 10.1101/2021.02.10.430657

**Authors:** Daniel C. Castro, Corinna S. Oswell, Eric T. Zhang, Christian E. Pedersen, Sean C. Piantadosi, Mark A. Rossi, Avery Hunker, Anthony Guglin, Jose A. Morón, Larry S. Zweifel, Garret D. Stuber, Michael R. Bruchas

## Abstract

Mu-opioid peptide receptor (MOPR) stimulation alters respiration, analgesia, and reward behavior, and can induce addiction and drug overdose. Despite its evident importance, the endogenous mechanisms for MOPR regulation of appetitive behavior have remained unknown. Here we report that endogenous MOPR regulation of appetitive behavior in mice acts through a specific dorsal raphe to nucleus accumbens projection. MOPR-mediated inhibition of raphe terminals is necessary and sufficient to determine appetitive behavioral state while select enkephalin-containing NAc ensembles are engaged prior to reward consumption, suggesting that local enkephalin release is the source of endogenous MOPR ligand. Selective modulation of NAc enkephalin neurons and CRISPR-Cas9-mediated disruption of enkephalin substantiate this finding. These results isolate a fundamental endogenous opioid circuit for state-dependent appetitive behavior and suggest alternative mechanisms for opiate modulation of reward.

---

For centuries opioids and their derivatives have been used as potent analgesics ^1^. Decades of neuroscience research has identified the mu opioid peptide receptor (MOPR) as the principal means through which most exogenous and endogenous opioids act to produce analgesia and pain relief. Importantly, while MOPR selective opioids have great therapeutic potential, they also induce multiple undesirable effects, such as inhibited gut motility, substance use disorder (addiction), and severe respiratory depression ^2–4^. Despite efforts to understand these and related opioid effects, there remains a fundamental lack of insight into the basic neurobiological, circuit and cellular mechanisms through which endogenous mu-opioids (peptide/receptor) act to regulate appetitive behavior.

MOPRs are found throughout the central and peripheral nervous systems. The effects of exogenous mu-agonism on reward-related behaviors are likely driven through mesocorticolimbic circuits ^5,6^. Of special interest is the MOPR-enriched nucleus accumbens medial shell (mNAcSh), a unique subregion within the striatum thought to coordinate cognitive, incentive motivational, affective, homeostatic, and sensory information to regulate motivated behaviors ^6–8^. mNAcSh is primarily composed of two endogenous opioid peptide producing medium spiny populations, preprodynorphin (*Pdyn*) and preproenkephalin (*Penk*). These propeptides cleave into potent, efficacious endogenous MOPR agonists ^9,10^. The prominent presence of MOPRs and these two endogenous neuropeptide ligands in mNAcSh suggest that the means and mechanisms to activate MOPRs are present within this subregion. Pharmacological evidence from the last 25 years supports this hypothesis, as exogenous action of MOPRs in mNAcSh has been shown to dramatically modulate a variety of behaviors ^11–14^.

Here we uncover a unique circuit for endogenous mu-opioid regulation of reward behavior using a series of complementary, high-resolution approaches (**Table S1**). We report a pivotal role for MOPR-expressing enkephalinergic lateral dorsal raphe inputs to NAc medial shell (LDRN^MOPR^-mNAcSh), that, when activated by retrogradely-released mNAcSh enkephalin (mNAcSh^Penk^), cause potentiation of appetitive behavior. These results provide foundational insight into how endogenous opioid peptides determine state-dependent appetitive behavior through actions on previously unrecognized receptors, cells and circuits to enhance both natural and opioid-mediated appetitive behavior.

First, we sought to determine whether MOPRs in mNAcSh are important for state-dependent modulation of appetitive behaviors. To test this, we used a within subjects voluntary sucrose consumption paradigm in which mice were tested *ad libitum* (baseline state), or after 24 hours of food deprivation (potentiated state) (Fig. 1a), along with a classic intracerebral microinjection approach to locally antagonize MOPR function in mNAcSh (Fig. 1b). Food deprived/vehicle mice significantly increased sucrose consumption relative to their *ad libitum*/vehicle test day. In contrast, MOPR antagonism in mNAcSh significantly reduced food deprived potentiation of sucrose consumption by ∼50%, but had no impact on sucrose intake on the *ad libitum* test day. Neither bilateral CTAP infusions into nearby brain regions nor unilateral mNAcSh hits reduced sucrose intake (Extended Data Fig. 1a-c).

**Fig. 1.**
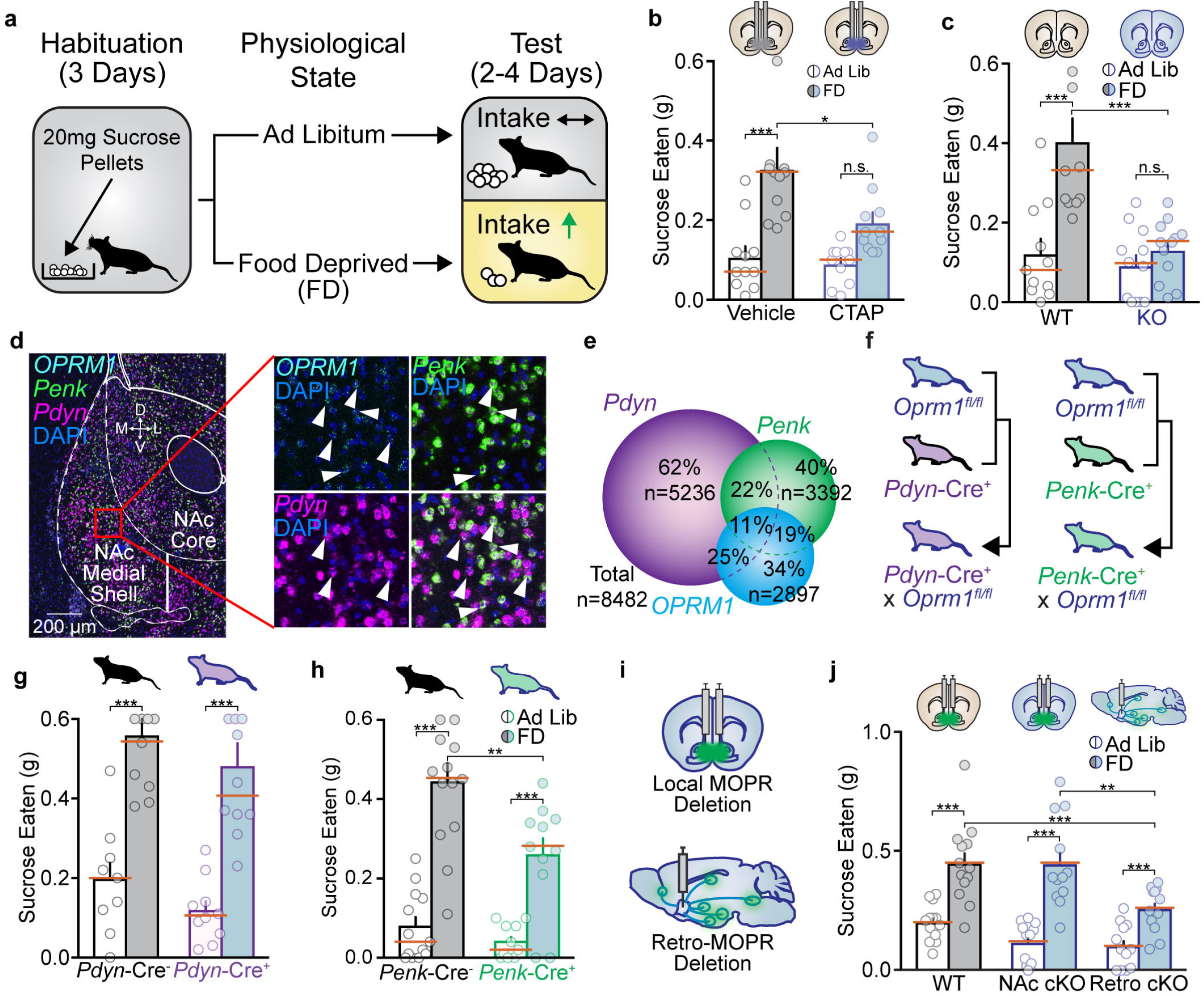
Endogenous MOPR activation in mNAcSh is necessary for potentiating state-dependent appetitive behavior. **a**. Schematic of voluntary sucrose consumption assay. **b**. CTAP reduces hunger enhanced, but not ad libitum, intake by 50% compared to within subjects vehicle (gray) control days (n = 11, 4 counter balanced test days, separated by 48 hours each; One-way ANOVA F = 15.84, p **<** 0.001; Ad Lib/vehicle vs FD/vehicle t(10) = 5.827, p **<** 0.001, Ad Lib/CTAP vs FD/CTAP t(10) = 3.012, p = 0.051, Ad Lib/vehicle vs Ad Lib/CTAP t(10) = 0.5273, p = 0977, FD/vehicle vs FD/CTAP t(10) = 3.667, p = 0.017; medians marked with orange bars). c. *Oprm1* KO show normal ad libitum intake, but 100% suppression of food deprived enhanced intake (n = 11WT, 12 KO, 2 counter balanced test days; Two-way ANOVA F(1,21) = 64.48, p **<** 0.001; Ad Lib/WT vs FD/WT t(21) = 12.91, p **<** 0.001, Ad Lib/KO vs FD/KO t(21) = 1.874, p = 0.144; Ad Lib/WT vs Ad Lib/KO t(42) = 0.5645, p = 0.82, FD/WT vs FD/KO t(42) = 5.107, p **<** 0.001). **d and e**. In situ hybridization **(d)** and quantification **(e)** of *Pdyn, Penk* and *Oprm1* in NAc medial shell (scale bar= 200µm). **f**. Schematic of *Oprm1* ^*fVfl*^ x *Pdyn-Cre* or *Penk-Cre* mouse line cross. **g**. Loss of MOPRs on *Pdyn-Cre+* neurons did not disrupt normal ad libitum or food deprived enhanced intake compared to *Pdyn-Cre-* littermate control mice (n = 9 cre-, 10 Cre+, 2 counter balanced test days; Two-way ANOVA F(1, 17) = 0.001, p = 0.972; Ad Lib/Cre-vs FD/Cre-t(17) = 8.808, p **<** 0.001, Ad Lib/Cre+ vs FD/Cre+ t(17) = 9.337, p **<** 0.001, Ad Lib/Cre-vs Ad Lib/Cre+ t(34) = 1.123, p = 0.466, FD/Cre-vs FD/Cre+ t(34) = 1.095, p = 0.483). h. Loss of MOPRs on *Penk-Cre+* neurons did not disrupt normal ad libitum intake, but suppressed food deprived enhanced intake by 50% compared to *Penk-Cre-* littermate control mice (n = 13 Cre-, 11 Cre+, 2 counter balanced test days; Two-way ANOVA F(1,22) = 4.434, p = 0.047; Ad Lib/Cre-vs FD/Cre-t(22) = 7.794, p < 0.001, Ad Lib/Cre+ vs FD/Cre+ t(22) = 4.308, p < 0.001, Ad Lib/Cre-vs Ad Lib/Cre+ t(44) = 0.7602, p = 0.699, FD/Cre-vs FD/Cre+ t(44) = 3.624, p = 0.001). i. Schematic local (top) or retrograde (bottom) MOPR deletion in NAc medial shell of *Oprm1* ^*fl/fl*^ mice. j. Loss of MOPRs in NAc did not reduce ad libitum or food deprived enhanced intake, whereas loss of mu receptors on neurons that project to medial shell suppressed food deprived enhanced food intake by 50% (n = 13 wildtype, 12 local NAc, 11 retro-NAc, 2 counter balanced test days; Two-way ANOVA F(2,34) = 6.046, p = 0.006; Ad Lib/WT vs FD/WT t(34) = 7.725, p **<** 0.001Ad Lib/NAc vs FD/NAc t(34) = 9.313, p < 0.001, Ad Lib/Retro vs FD/Retro t(34) = 4.398, p < 0.001, Ad Lib/WT vs Ad Lib/NAc t(68) = 1. 730, p = 0.242, Ad Lib/WT vs Ad Lib/Retro t(68) = 1.997, p = 0.142, Ad Lib/NAc vs Ad Lib/Retro t(68) = 0.2616, p = 0.991, FD/WT vs FD/NAc t(68) = 0.08336, p = 0.999, FD/WT vs FD/Retro t(68) = 3.834, p **<** 0.001, FD/NAc vs FD/Retro t(68) = 3.678, p = 0.001).

To determine the ultimate extent to which MOPRs can enhance appetitive intake, we tested wildtype and genetically modified *Oprm1* constitutive knockout mice in the same voluntary sucrose consumption paradigm (Fig. 1c). Loss of MOPR function in *Oprm1* KO mice did not disrupt *ad libitum* sucrose intake. In contrast, food deprivation in *Oprm1* KO mice completely blocked potentiated sucrose consumption. These data indicate that MOPRs in mNAcSh account for ∼50% of all MOPR-mediated potentiation of sucrose consumption, indicating that this is critical brain site for MOPR regulation of appetitive behavior.

We next determined if MOPRs were preferentially expressed on a specific cellular subtype using fluorescent *in situ* hybridization (FISH). We found that MOPRs are expressed on ∼30-35% of all neurons within mNAcSh, and on about ∼50% of *Pdyn* or *Penk* subpopulations (Extended Data Fig. 1d-e). To test whether MOPRs preferentially act on one subpopulation to potentiate sucrose consumption, we crossed MOPR conditional knockout mice (*Oprm1*^*fl/fl*^) with *Pdyn*-Cre or *Penk*-Cre mouse lines to selectively delete MOPRs from each respective cell-type (Fig. 1f, Extended Data Fig. 1d-g). We found that loss of MOPRs on *Penk*-Cre^+^ but not *Pdyn*-cre^+^ MSNs resulted in a 50% reduction in food deprived potentiation of sucrose consumption (Fig. 1g-h). This cell-type specific effect appeared to selectively drive appetitive behaviors, as loss of MOPRs on *Penk*-Cre^+^ cells did not disrupt normal avoidance of the open arms in an elevated zero maze after restraint stress, even though systemic naloxone injections and *Oprm1* KO mice did disrupt avoidance behaviors. These results suggest that MOPR-dependent appetitive and avoidance behaviors are mediated by separate subsystems (Extended Data Fig. 2).

MOPR deletions on *Penk*-Cre^+^ neurons left unresolved whether the reduction in food deprived potentiation of sucrose consumption is mediated by accumbens or non-accumbens cells. Therefore, we conditionally deleted MOPRs in mNAcSh of *Oprm1*^*fl/fl*^ mice (Extended Data Fig. 1h-i). Local deletion did not reduce sucrose intake relative to wildtype controls (Fig. 1j). Since local pharmacological antagonism did reduce intake, we reasoned that MOPRs may act presynaptically. We directly tested this by injecting retrograde viruses into mNAcSh of *Oprm1*^*fl/fl*^ mice to delete MOPRs on neurons that project to medial shell ^15^ (Extended Data Fig. 1j). We found that retrograde MOPR deletion reduced potentiated sucrose consumption by ∼50% (Fig. 1j). Collectively, these data indicate that presynaptic MOPRs in mNAcSh are recruited in a state-dependent manner to potentiate ongoing appetitive behavior, and are located on a *Penk*^+^ neuronal population.

We next used a retrograde viral tracing approach to identify specific enkephalinergic projections to mNAcSh (Fig. 2a, Extended Data Fig. 3a). Fluorescent labeling revealed several brain regions, including an unrecognized lateralized population of dorsal raphe nucleus (LDRN). Functional experiments in other labeled sites failed to modulate food deprived intake (Extended Data Fig. 4). Therefore, we focused efforts on understanding MOPRs role within the LDRN population. Using FISH, we found that DRN cells expressed *Penk* (28%), *Oprm1* (31%), and *Tph2* (19%). About half of DRN^Penk^ neurons coexpressed *Oprm1* (14%), but only a very small subset additionally expressed *Tph2* (4%) (Fig. 2b-c). Combinatorial viral tracing and FISH experiments confirmed high coexpression of *Penk* and *Oprm1* in neurons that specifically project to mNAcSh (60% overlap) (Extended Data Fig. 2d-j). Considering the high degree of overlap, we will refer to this specific projection as LDRN^MOPR^-mNAcSh.

**Fig. 2.**
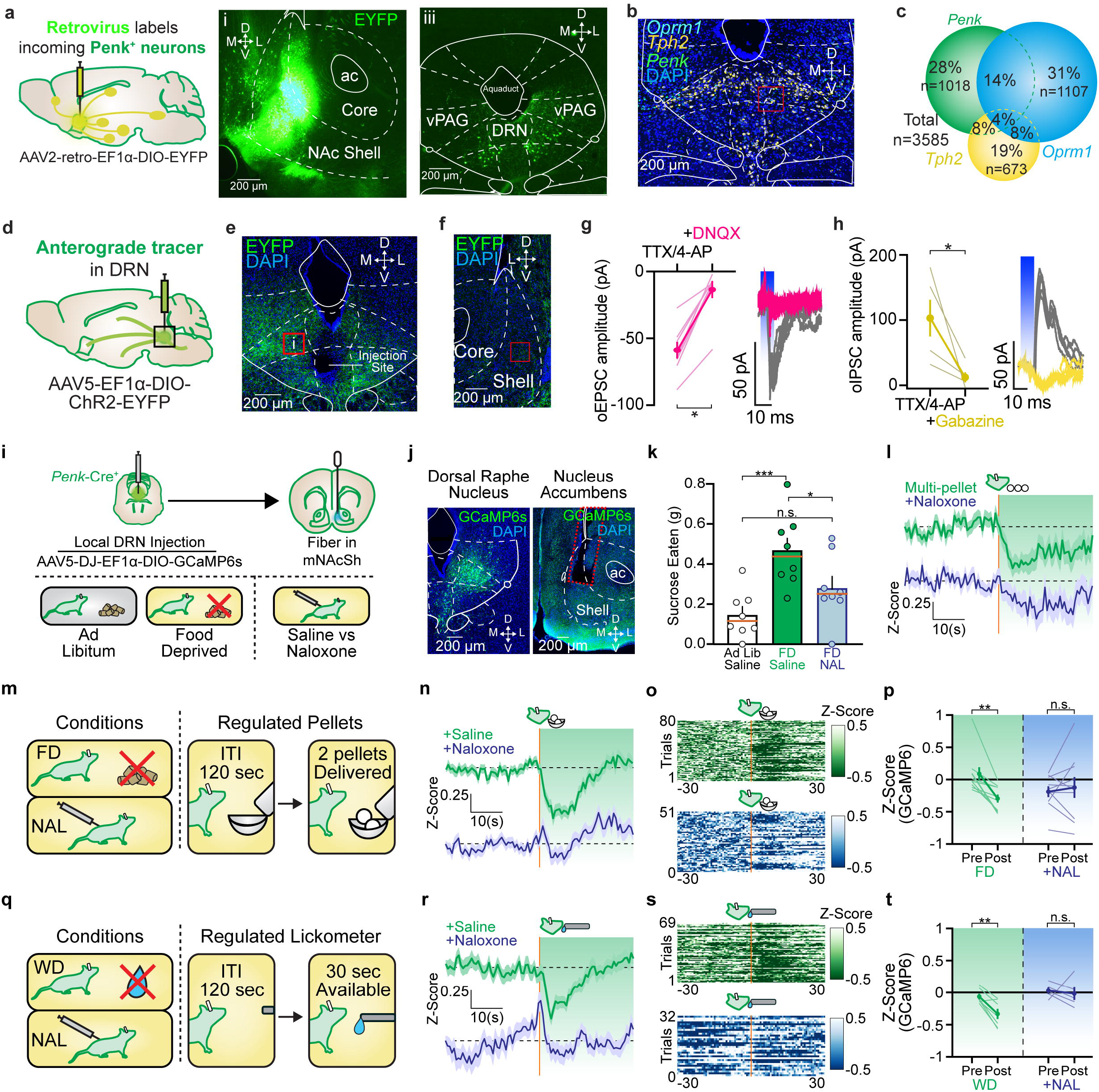
LDRN^Penk^-NAc projections make monosynaptic connections and display opioid-dependent spatiotemporal signaling dynamics. **a**. Schematic (left) of a retrograde fluorescently tagged virus injection into medial shell of a *Penk-Cre+* mouse (middle, scale bar = 200 µm) and labeled neurons in dorsal raphe nucleus (right). b. Fluorescent in situ hybridization of *Oprm1, Tph2* and *Penk* in dorsal raphe nucleus (scale bar = 200 µm). **c**. Quantification of FISH from panel B. **d and e**. Schematic and image of a local injection of AAV5-Ef1a-DIO-EYFP into DRN^**Penk**^ (scale bar = 200 µm). **f**. Image of dorsal raphe projection fibers from e (scale bar = 200 µm). **g**. oEPSC amplitude was reduced by the application of DNQX (pink, t-test, t(7) = 8.48, p < 0.001). Blue shaded region indicates duration of optical stimulation. **h**. olPSC amplitude was reduced by the application of gabazine (yellow, paired I-test, t(4) = 3.22, p = 0.03). i. Schematic of fiber photometry and voluntary sucrose consumption paradigm. **j**. Expression of GCaMP6s in DRN^**Penk**^ (left, scale bar = 200 µm) and fiber placement in mNAcSh (scale bar = 200 µm). k. Food deprivation increases food intake (green, FD) relative to ad libitum intake (white, Ad Lib), and is reduced by systemic naloxone (blue, NAL) (n = 8, 3 counter balanced test days; Two-Way ANOVA F(2,7) = 3.032, p <0.001; Ad Lib/saline vs FD/saline t(14) = 5.253, p < 0.001, Ad Lib/saline vs FD/naloxone t(14) = 2.204, p = 0.128, FD/saline vs FD/naloxone t(14) = 3.049, p = 0.026). I. Average Z-scored trace aligned to onset of multi-pellet consumption of LDRN^**Penk**^-mNAcSh. FD/saline trace (green) shows rapid and sustained inhibition. FD/naloxone trace (blue) shows blunted response. **m**. Schematic of regulated food intake paradigm. Two pellets were non-contingently delivered every 90-120 seconds for 30 minutes. Mice were tested in either FD/saline or FD/naloxone conditions. n. Average Z-scored trace aligned to onset of multi-pellet consumption of LDRN^**Penk**^-mNAcSh. Food deprived/saline trace (green) shows rapid and sustained inhibition. Food deprived/naloxone trace (blue) shows blunted response. **o**. Heatmap of individual trials across all tested mice in food deprived/saline (green) or food deprived/naloxone test days. Orange lines indicate onset of pellet consumption. p. Quantification of the average Z-score twenty seconds prior to the onset of multi-pellet pellet bouts versus twenty seconds after the onset. FD/saline traces (green) show significant reductions in GCaMP6s activity whereas FD/naloxone traces (blue) do not (FD/saline/pre vs FD/saline/post t(9) = 4.360, p = 0.002, FD/naloxone/pre vs FD/naloxone/post t(9) = 0.5361, p = 0.605). q. Schematic of regulated lickometer paradigm. The lickometer was non-contingently presented every 90-120 seconds for 30 seconds. Mice were tested in either WD/saline or WD/naloxone conditions. WD = Water Deprived. r. Average Z-scored trace aligned to onset of sustained licking bouts (longer than 2 seconds) of LDRN^**Penk**^-mNAcSh. WD/saline trace (green) shows rapid and sustained inhibition. WD/naloxone trace (blue) shows blunted response. **s**. Heatmap of individual trials across all tested mice in WD/saline (green) or WD/naloxone test days. Orange lines indicate onset of licking bout. **t**. Quantification of the average Z-score twenty seconds prior to the onset of sustained licking bouts versus twenty seconds after the onset. WD/saline traces (green) show significant reductions in GCaMP6s activity whereas WD/naloxone traces (blue) do not. Some mice did not lick on naloxone treated days and were therefore not included in this analysis (WD/saline/pre vs WD/saline/post t(6) = 4.444, p = 0.004, WD/naloxone/pre vs WD/naloxone/post t(4) = 0.3639, p = 0.734).

Finally, we determined whether LDRN^MOPR^-mNAcSh projections make functional monosynaptic connections with mNAcSh using optogenetics and *ex vivo* patch-clamp electrophysiology. Channelrhodopsin labeled LDRN^MOPR^-mNAcSh inputs were observed in the central portion of mNAcSh (Fig. 2d-f). Photo-stimulation of LDRN terminals revealed that about 15% of NAc neurons received monosynaptic connections. Of these monosynaptically identified cells, ∼7% elicited DNQX-sensitive EPSPs, ∼3% elicited Gabazine-sensitive IPSPs, and ∼4% showed both (Fig. 2g-h). Taken together, these results indicate that a MOPR-expressing LDRN-mNAcSh circuit that is distinct from more canonical opioid/serotonin systems, could represent a critical locus for endogenous opioid modulation of appetitive behavior ^16–19^.

We next determined whether the *in vivo* activity of the LDRN^MOPR^-mNAcSh projection was 1) related to time-locked appetitive behaviors, and 2) are opioid dependent. First we expressed the calcium indicator GCaMP6s into LDRN of *Penk*-Cre^+^ mice ^20^ and implanted an optical fiber into mNAcSh to measure changes in terminal fluorescent activity. We then tested the mice on the voluntary sucrose paradigm after systemic saline or naloxone injections (Fig. 2i-j). We found that compared to the infrequent *ad libitum* consummatory behavior, food deprived mice engaged in sustained eating bouts whereby they ate multiple pellets in a row (i.e., multi-pellet bout, Fig. 2k, Extended Data Fig. 5a). MOPR antagonism with naloxone pretreatment reduced multi-pellet bout behavior, suggesting this phenotype was state and endogenous opioid dependent. When we aligned LDRN^MOPR^-mNAcSh GCaMP activity to the onset of multi-pellet bouts, we observed a sustained inhibition of GCaMP activity (∼20 seconds) coinciding with the total duration of the eating bout (Fig. 2l, Extended Data Fig. 5b-d). The reduction of GCaMP activity on LDRN^MOPR^-mNAcSh terminals during multi-pellet bout consummatory events was attenuated by the opioid antagonist naloxone.

Due to inherent variability of consummatory behavior within and between freely feeding mice, we also utilized a regulated sucrose model to standardize pellet consumption (**Fig. 2m**). We observed an immediate and sustained reduction in GCaMP activity during consumption of sucrose on LDRN^MOPR^-mNAcSh terminals (Fig. 2n-o). Additionally, terminal inhibition and overall pellet consumption was attenuated by naloxone pretreatment (Extended Data Fig. 5e). Quantification of the average of the trace (Z-score) prior to versus after the onset of a multi-pellet bout significantly confirmed the reduction in GCaMP activity; this reduction was absent following naloxone pretreatment (Fig. 2p).

Next, we determined whether LDRN^MOPR^-mNAcSh terminals activity would be similarly inhibited during liquid reward consumption by modifying the food deprivation/sucrose pellet model into a water deprivation/sucrose solution model (Fig. 2q). During this test, mice received non-contingent, intermittent access to a sucrose solution via lickometer to induce lick bouts (similar to multi-pellet bouts). Here we observed immediate and sustained inhibition of LDRN^MOPR^-mNAcSh terminals when mice initiated licking bouts following water deprivation, which was similarly blocked by opioid receptor antagonist naloxone (Fig.2r-t). These results indicate that the temporally precise recruitment of a previously unrecognized endogenous opioid circuit acts to potentiate appetitive behavior.

To test whether MOPRs on the LDRN-mNAcSh projection causally mediate the potentiation of appetitive behaviors, we selectively restored MOPR within LDRN^Penk^ neurons in *Oprm1* KO x *Penk*-Cre^+^ mice via targeted viral injection (Fig. 3a-b, Extended Data Fig. 6a). MOPR gain of function in only LDRN^Penk^ neurons significantly restored food deprived potentiation of sucrose intake (Fig. 3c). Similarly, MOPR rescue in LDRN^Penk^ neurons partially and significantly restored morphine conditioned place preference (Fig. 3d). However, MOPR rescue in this circuit was unable to restore morphine analgesia. This indicates that LDRN^Penk^ neurons are critical for modulating both endogenous and exogenous opioid reward-related behaviors, but do not mediate analgesia.

**Fig. 3.**
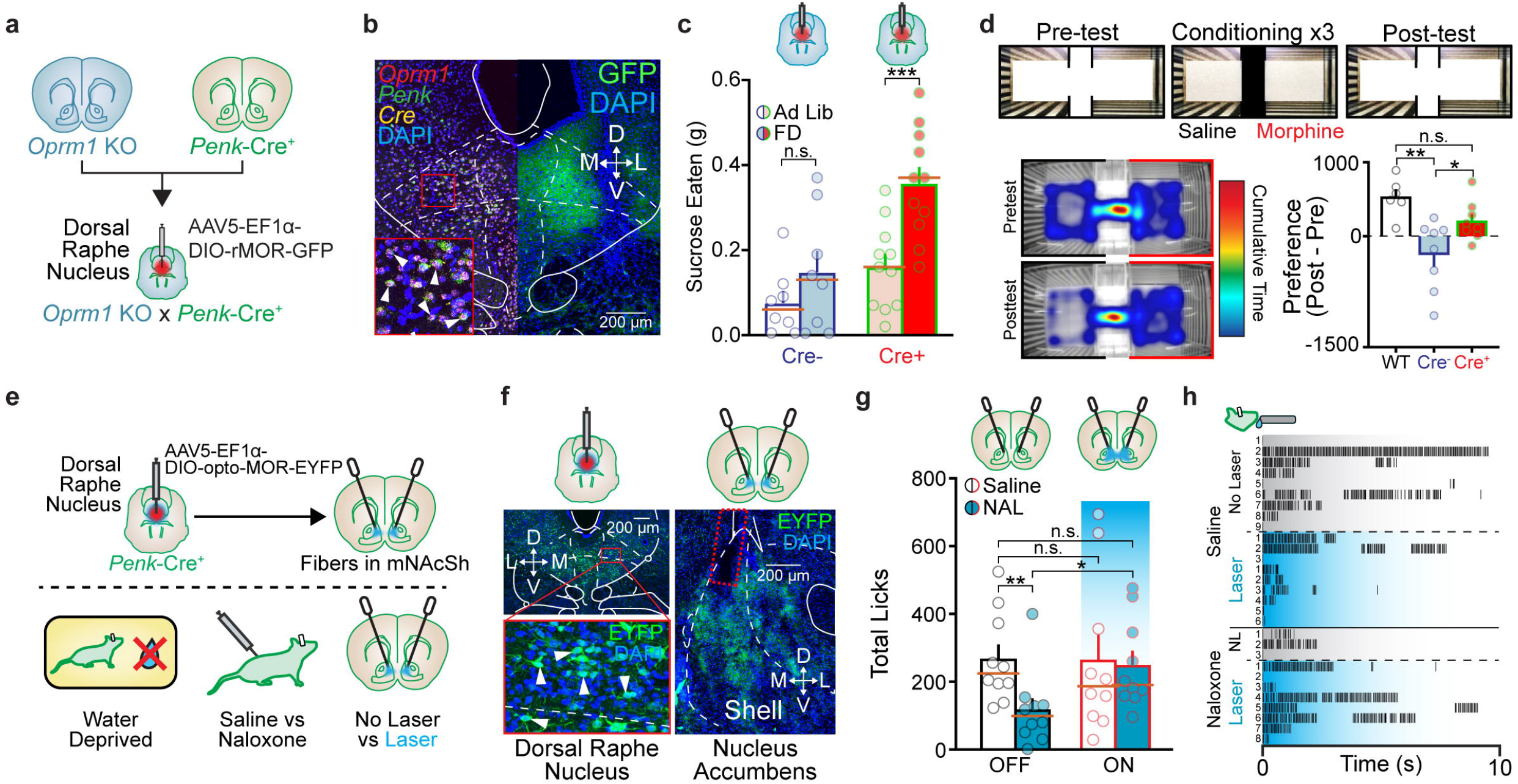
MOPR activation on LDRN MOPR.m NAcSh is sufficient to potentiate appetitive behavior. **a**. Schematic of *Oprm1* KO x *Penk*-Cre rescue of MOPRs in IDRN. b. In situ micrograph (left; *Oprm1, Penk*, Cre) and associated fluorescent micrograph (right; GFP-tagged viral vector) of AAV5-Ef1a-DIO-rMOR-EYFP in DRN of *Oprm1* KO x *Penk*-Cre+ mice (scale bar = 200 µm). c. *Oprm1* KO x *Penk-Cre-* mice show no increase in food intake after food deprivation. *Oprm1* KO x *Penk-Cre+* mice how robust enhancement of intake after food deprivation mice (n = 8 Cre-, 11 Cre+, 2 counter balanced test days; Two-way ANOVA F(1, 17) = 6.926, p = 0.017; Cre-ad libitum vs Cre-deprived t(17) = 2.025, p = 0.114, Cre+ ad libitum vs Cre+ deprived t(17) = 6.430, p < 0.001, Cre-ad libitum vs Cre+ ad libitum t(34) = 1.603, p = 0.223, Cre-deprived vs Cre+ deprived t(34) = 3.913, p < 0.001). d. Morphine conditioned place preference (CPP) assay. (top) Schematic of CPP procedure. (bottom left) Heat map of cumulative time spent in each location before (“pretest”) and after (“posttest”) morphine conditioning. Warmer colors indicate more time spent in that area. (bottom right) Wildtype and *Oprm1* KO x *Penk*-Cre+ mice spent significantly more time in the morphine paired side compared to *Oprm1* KO x *Penk-Cre-* mice (n = 6 WT, 8 *Oprm1* KO x *Penk-Cre-*, 8 *Oprm1* KO x *Penk*-Cre+; One-way ANOVA F(2, 19) = 8.684, p = 0.002; WT Preference vs Cre-Preference t(19) = 5.79, p = 0.002, WT Preference vs Cre+ Preference q(19) = 2.403, p = 0.231, Cre+ Preference vs Cre-Preference t(19) = 3.658, p = 0.046). e. Schematic of opto-MOR experiments. f. Expression of EYFP-tagged opto-MOR in LDRN ^**Penk**^ (left, scale bar = 200 µm) and fiber placement (red dashed line) in mNAcSh (right, scale bar = 200 µm). Zoomed in image shows labeled nuclei in DRN (bottom left, red box). g. Saline/No Laser treated mice licked significantly more for a sucrose solution compared to the naloxone/No Laser treated test day. Opto-MOR stimulation had no effect on intake after saline, but restored licking after naloxone mediated suppression (n = 8, 4 counter balanced test days, separated by 48 hours each; One-way ANOVA F(1.743, 12.2) = 6.485, p = 0.014; saline/No Laser vs naloxone/No Laser t(7) = 4.838, p = 0.011, saline/No Laser vs saline/Laser t(7) = 0.4549, p = 0.663, saline/No Laser vs naloxone/Laser t(7) = 2.131, p = 0.197, saline/Laser vs naloxone/No Laser t(7) = 3.328, p = 0.05, saline/Laser vs naloxone/Laser t(7) = 1.451, p = 0.344, naloxone/No Laser vs naloxone/Laser t(7) = 3.494, p = 0.049). Medians marked in orange. h. Raster plots of individual licking events for one mouse, separated by trial. Only trials in which mice licked were included. Naloxone injections reduced number and length of lick bouts, which was reversed by opto-MOR stimulation.

We next determined whether spatiotemporal engagement of MOPR signaling on LDRN^MOPR^-mNAcSh terminals controls potentiation of sucrose consumption in a time-locked manner. We expressed the light activated chimeric receptor opto-MOR on LDRN^Penk^ neurons and implanted fibers into mNAcSh for terminal specific spatially localized activation of MOPR signaling (Fig. 3e-f) ^21^. We then trained the mice on the intermittent sucrose solution task (Fig. 3k), and tested whether selective stimulation of presynaptic MOPR signaling via opto-MOR on LDRN^Penk^-mNAcSh terminals could circumvent naloxone mediated opioid antagonism. We found that naloxone alone reduced overall licks compared to the saline treated day (Fig. 3g). However, photo-activation of opto-MOR after naloxone treatment was sufficient to significantly restore licking behavior back to baseline levels. Further analysis revealed that behaviorally initiated and time-locked opto-MOR stimulation restored the sustained licking behavior that was blunted by naloxone (Fig. 3h). An opposing experiment in which we artificially stimulated LDRN^Penk^-mNAcSh terminals via ChR2 during lick behavior demonstrated that inhibition of this pathway is necessary for deprivation potentiation of sucrose intake (Extended Data Fig. 6d-h). Taken together with pharmacological blockade and genetic deletion of MOPR (Fig 1b,j), these findings indicate that time locked, MOPR-mediated inhibition of the LDRN^Penk^-mNAcSh circuit is both necessary and sufficient for potentiating appetitive behaviors.

Since mNAcSh enkephalin neurons make up nearly 50% of all neurons within the NAc, we hypothesized that this population could activate presynaptic MOPRs on LDRN^Penk^-mNAcSh terminals via retrograde neuropeptide release and action. To determine whether this population is meaningfully recruited during consummatory behavior, we expressed GCaMP6s in mNAcSh^Penk^ neurons and implanted a GRIN lens for endoscopic single-cell calcium imaging during voluntary sucrose intake (Fig. 4a-c, Extended Data Fig. 7a). We reliably tracked single-cell calcium activity in 143 of the same neurons within mNAcSh across both *ad libitum* and food deprived test days (Fig. 4d-f). On the food deprived test day, we observed a mean response from all cells of mild inhibition at the onset of multi-pellet bout consumption (Fig. 4g) as has been previously suggested using electrophysiology ^22,23^. However, this genetically isolated single cell tracking approach revealed that mNAcSh^Penk^ neurons were not uniformly inhibited, and instead, showed one of several distinct response patterns (Fig. 4f, Extended Data Fig. 7b-e). *K-means* clustering isolated four unique subpopulations of neurons (clusters), which included: **1)** Pre-onset activated (4%), **2)** Onset activated (8%), **3)** Onset inhibited (19%), and **4)** Non-responsive (69%) (Fig. 4g-l). These clusters were also separable from clusters that were responsive to sucrose sniffing, rearing, or grooming (Fig. Extended Data Fig. 7f-i). These results suggest that behaviorally identified local mNAcSh^Penk^ neuronal populations are selectively recruited to enhance appetitive behavior and may act to provide the endogenous source of MOPR ligand acting on the LDRN^MOPR^-mNAcSh^Penk^ circuit.

**Fig. 4.**
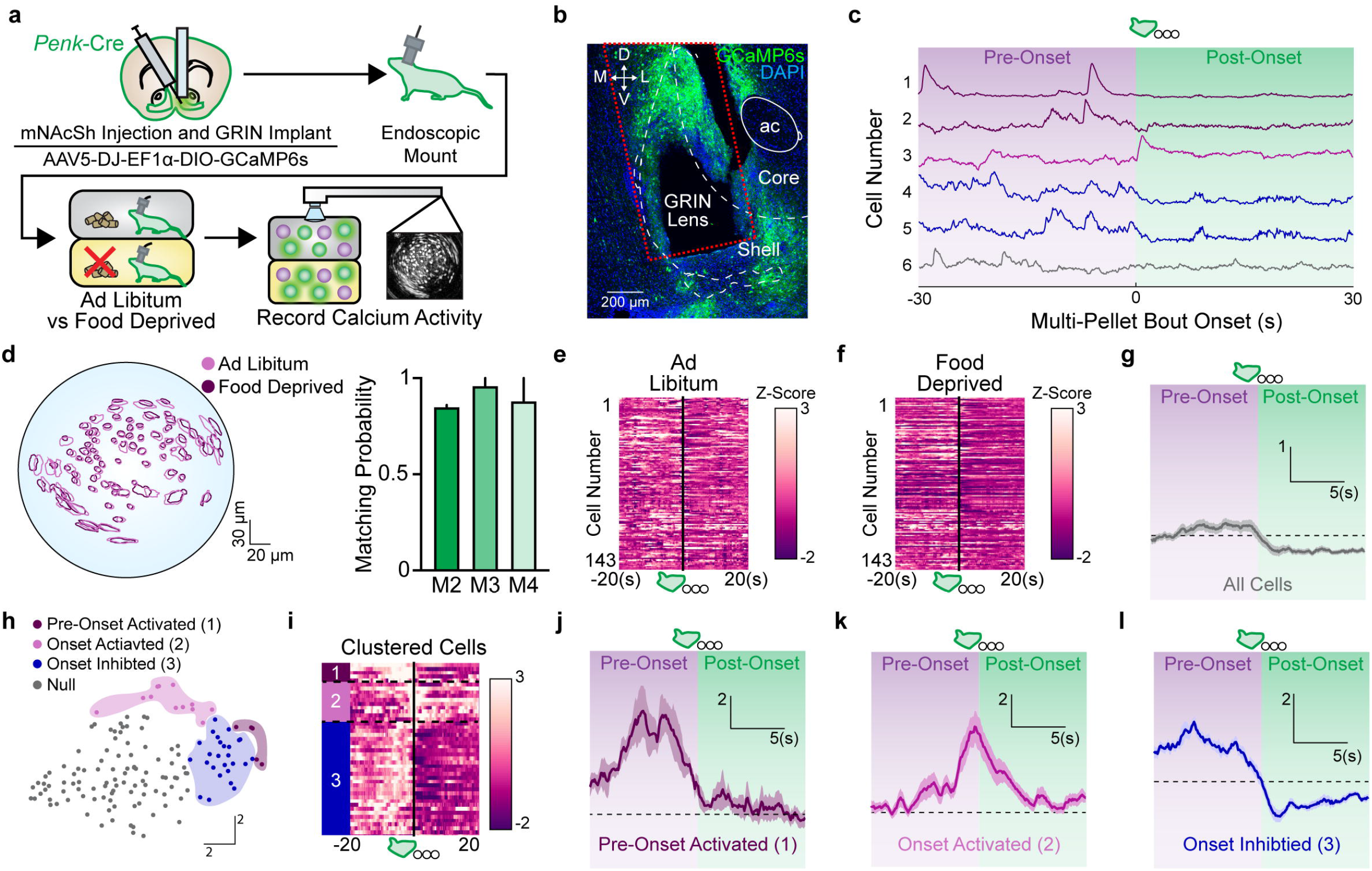
mNAcSh enkephalinergic ensembles are modulated by physiological state and potentiate appetitive behavior. **a**. Schematic of *in vivo* 1-photon imaging experiments. **b**. Expression of AAV-DJ-EF1a-DIO-GCaMP6s and GRIN lens placement (red dashed line) in mNAcSh of *Penk-Cre+* mouse (scale bar = 200 µm). c. Examples of individual cell traces aligned to initiation of a multi-pellet bout. **d**. Example cell map (left) for cells that were matched (right) across ad libitum and food deprived test days. **e and f.** Heat plot of all 143 matched cells on the ad libitum or food deprived test day. Change in Z-scored fluorescence displayed by greater signal (white) or lesser signal (purple). Cells aligned to onset of multi-pellet bouts. **g**. Average trace (left) of all tracked cells. **h**. Tsne plot color coded to show four distinct clusters of neurons during multi-pellet bout consumption, determined using kmeans clustering analysis. **i.** Heat plot of the three behaviorally modulated clusters, each cluster separated by black dashed lines and color coded to match H. Change in Z-scored fluorescence displayed by greater signal (white) or lesser signal (purple). Cells aligned to onset of multi-pellet bouts. **j**. Average trace of Pre-onset activated neurons (cluster 1). **k**. Average trace of Onset activated neurons (cluster 2). I. Average trace of Onset inhibited neurons (cluster 3).

To test the necessity of local mNAcSh^Penk^ in potentiating appetitive consumption, we induced cell-type specific ablations of mNAcSh^Penk^ neurons via caspase viral injections (Fig. **5a-c**), thereby eliminating this neuronal population and corresponding production of the endogenous opioid neuropeptide enkephalin. Mice were then tested in the voluntary sucrose consumption paradigm. Caspase ablation of mNAcSh^Penk^ neurons significantly reduced food deprived sucrose consumption, but had no impact on *ad libitum* intake (Fig. 5d). FISH experiments further established that we effectively ablated mNAcSh^Penk^ neurons (Fig. 5b-c).

**Fig. 5.**
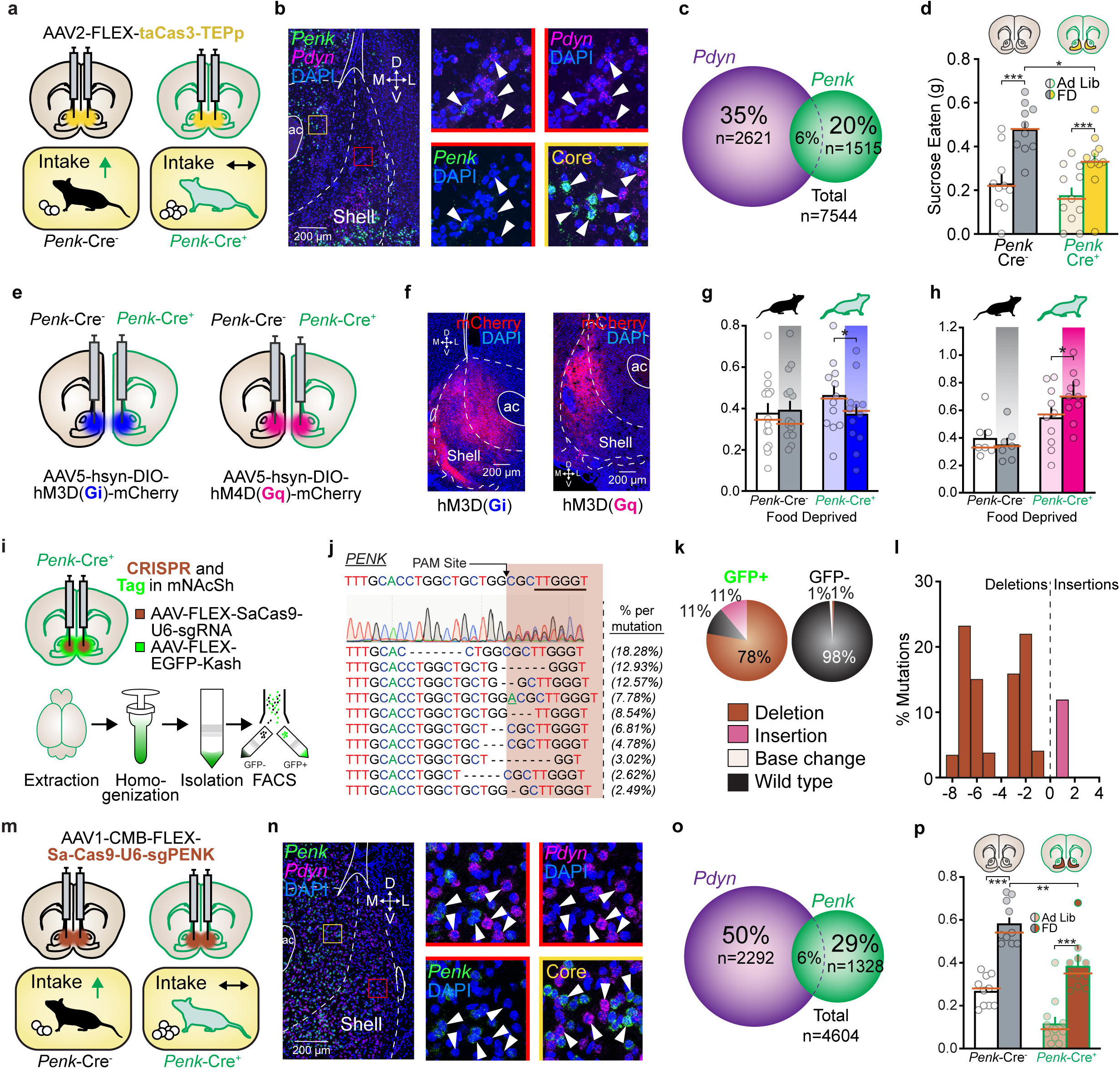
Endogenous mu-opioid peptide within mNAcSh is necessary for potentiating appetitive behavior. **a**. Schematic of food intake paradigm after caspase-mediated ablation of mNAcSh^Penk^. **b and c**. In situ hybridization (b, scale bar= 200 µm) and quantification (c) of *Pdyn* and *Penk* in mNAcSh after caspase injections. **d.** Caspase injections in mNAcSh did not disrupt ad libitum or food deprived intake in *Penk-Cre-* mice, but did blunt expected food deprived enhancement in *Penk-Cre+* mice (n = 10 Cre-, 11 Cre+, 2 counter balanced test days; Two-way ANOVA F(1, 19) = 3.994, p = 0.06; Cre-ad libitum vs Cre-deprived t(19) = 7.399, p < 0.001, Cre+ ad libitum vs Cre+ deprived t(19) = 4.865, p < 0.001, Cre-deprived vs Cre+ deprived t(38) = 2.553, p = 0.029). **e.** Schematic of hM3D(Gi) (left) and hM3D(Gq) (right) DREADD experiments. **f.** Fluorescent micrograph of mCherry-tagged enkephalin cells in mNAcSh for hM3D(Gi) (left) and hM3D(Gq) (right) experiments (scale bar= 200µm). **g.** CNO injections suppressed hunger enhanced intake in Cre+ mice, but had no effect in Cre-mice (n = 15 Cre-, 12 Cre+, 4 counter balanced test days; One-way ANOVA Cre-F(14,42) = 5.688, p < 0.001; Cre-deprived/saline vs Cre-deprived/CNO t(14) = 0.4408, p = 0.889, Cre+ deprived/saline vs Cre+ deprived/CNO t(11) = 2.9, p = 0.029). **h.** CNO injections increased intake above the already elevated food deprived intake in Penk-Cre+ mice, but had no effect in Penk-Cre-mice (Two-Way ANOVA F(3,45) = 5.815, p = 0.002; Cre-FD/saline vs Cre-FD/CNO t(6) = 2.22, p = 0.346, Cre+ FD/saline vs Cre+ FD/CNO t(9) = 3.492, p = 0.04). **i.** Schematic of CRISPR virus development and validation. **j.** Sequencing of GFP+ nuclei: (Top) sgPenk sequence with PAM underlined and SaCas9 cut site indicated by black arrow. (Middle) Sanger sequencing results displaying multiple peaks beginning at the SaCas9 predicted cut site. (Bottom) Top ten mutations at cut site with the percent of occurrence on the left. Insertions: underlined. Deletions: marked with”-”. Affected sites after SaCas9 insertion: shaded brown. **k.** Percent of wild type (black), deletions (brown), insertions (pink), and base changes (white) as percent of total reads for GFP+ and GFP-nuclei. **I.** Frequency distribution of insertions (pink) and deletions (brown) for *Penk* from GFP+ nuclei. **m.** Schematic of food intake paradigm after CRISPR-mediated disruption of enkephalin production in mNAcSh. **n and o**. In situ hybridization (n, scale bar= 200 µm) and quantification (o) of *Pdyn* and *Penk* in mNAcSh after CRISPR viral injections. **p**. CRIS PR virus injections in mNAcSh shell did not disrupt ad libitum or food deprived intake in *Penk-Cre-* mice, but did blunt expected ad libitum and food deprived enhancement in *Penk-Cre+* mice (n = 10 Cre-, 11 Cre+, 2 counter balanced test days; Two-way ANOVA F(1, 19) = 2.501, p = 0.13; Cre-ad libitum vs Cre-deprived t(19) = 14.7, p < 0.001, Cre+ ad libitum vs Cre+ deprived t(19) = 13.13, p < 0.001, Cre-ad libitum vs Cre+ ad libitum t(38) = 3.396, p = 0.003, Cre-deprived vs Cre+ deprived t(38) = 4.444, p < 0.001).

We next sought to determine whether modulation, rather than ablation, of local mNAcSh^Penk^ neurons could bidirectionally alter voluntary sucrose consumption. To test the necessity of this population, we expressed the inhibitory DREADD HM4D(Gi) to inhibit neurons during the sucrose consumption tests after pretreatment with clozapine-N-oxide (CNO, Fig. 5e-f). Gi-DREADD activation reduced both *ad libitum* and food deprived sucrose consumption below their saline treated test days (Fig. 5g, Extended Data Fig. 8a). To test the sufficiency of mNAcSh^Penk^ neurons, we expressed the excitatory DREADD HM3D(Gq) and tested them in the voluntary sucrose task (Fig. 5i-j). CNO administration had no effect on sucrose intake in *the ad libitum* condition compared to the saline treated test day (Extended Data Fig. 8b). However, we did observe an increase in consumption above the potentiated food deprived sucrose intake, suggesting that Gq-DREADD stimulation can further augment the already engaged mNAcSh^Penk^ system (Fig. 5h). As a potential alternative, we also tested whether bidirectional DREADD modulation of POMC neurons (producers of the potent MOPR ligand beta-endorphin) in arcuate nucleus would affect sucrose intake. Neither Gi-nor Gq DREADD activation altered food deprived sucrose consumption (Extended Data Fig. 8c-g). These results indicate that mNAcSh^Penk^ neurons are the likely endogenous opioid source for potentiating appetitive behaviors.

To directly assess whether the local mNAcSh^Penk^ neuropeptide itself, rather than just *Penk*-expressing neurons, is recruited to potentiate appetitive behavior in the voluntary sucrose consumption task, we developed and used a single conditional CRISPR-mediated neuropeptide viral deletion strategy. First, we designed and packaged a single-viral SaCas9 guided mRNA construct ^24,25^ to selectively disrupt enkephalin neuropeptide production in a cell-type selective manner (Fig. 5i). FACS and PCR analyses indicated that this virus disrupts enkephalin mRNA production by 90%, primarily via base pair deletions, though base pair insertions also occurred (Fig. 5j-l, Extended Data Fig. 9a). When injected into mNAcSh, FISH experiments showed 30-50% reductions in *Penk*, with only the most ventral and lateral portions of the shell spared, likely due to the limitations of viral spread (Fig. 5m-o). Behaviorally, *Penk*-Cre^+^ mice showed a significant (∼50%) reduction in food deprived potentiation of sucrose consumption compared to *Penk*-Cre-mice injected with the same cre-dependent vector (Fig. 5p). *Penk*-Cre^+^ mice with only unilateral CRISPR infection did not display deficits in sucrose consumption (Extended Data Fig. 9b). Lastly, we also examined a potential role for local *Pdyn* ^26^and by default *Leu-Enk* by locally deleting it mNAcSh of *Pdyn*^*fl/fl*^ mice (Extended Data Fig. 9c). This peptide deletion (analogous to our *Penk* CRISPR disruption) did not show deficits in food deprived potentiation of sucrose consumption. Together, these results indicate that enkephalin peptide production in mNAcSh is specifically necessary for potentiating appetitive behavior, identifying a critical locus for endogenous opioid peptide action on reward seeking.

Here we isolated an endogenous mechanism underlying mu-opioid receptor modulation of appetitive behaviors in nucleus accumbens medial shell. A critical function for MOPRs within mNAcSh appears to be its role in mediating state-dependent potentiation of appetitive behavior. Across the many behavioral and MOPR/*Penk* selective manipulations we tested, the LDRN^MOPR^-mNAcSh^Penk^ circuit regulated 50% of all of MOPRs effects on appetitive behavior. This leaves unresolved: where are the remaining endogenous MOPRs and opioid peptides that regulate appetitive behaviors? One possibility is ventral tegmental area, within which mu-opioid stimulation has been shown to modulate phasic dopamine neuron activity and subsequent appetitive behaviors ^21,27,28^, yet a local source of endogenous opioid peptide in the region has yet to be established. Other potential key sites of endogenous MOPR action include the orbitofrontal cortex, dorsal striatum, basolateral amygdala and parabrachial nucleus ^6,29–32^. One difficulty in dissociating specific MOPR functions within selected brain areas/circuits is related to the type of behavior being investigated. For example, we showed here that potentiated avoidance in the elevated zero maze after restraint was blocked by either deleting MOPRs in *Oprm1* KO mice or by pretreatment with naloxone (Extended Data Fig. 2). However, *Oprm1*^*fl/fl*^ x *Penk*-Cre^+^ mice did not show such deficits, suggesting that an alternative “MOPR avoidance” circuit may exist in parallel. We also show that MOPRs on LDRN^Penk^ regulate appetitive sucrose intake, but not morphine analgesia, indicating a possible third “MOPR analgesia” circuit. Such demonstrations for non-appetitive MOPR functions have been reported and would be consistent with other endogenous neuropeptide systems ^21,33–39^. For example, the dynorphin/kappa opioid receptor system, which has canonically been ascribed “aversive” roles, has now been shown to oppositely enhance appetitive or positive affective behaviors depending on its activation within dorsoventral or rostrocaudal sites in mNAcSh ^13,40,41^. Similarly, NOPR stimulation in para-nigral VTA intensely suppresses responding for sucrose rewards, whereas stimulation in mNAcSh oppositely increases food intake ^39,42^.Differences in homogenous vs heterologous state-dependent neuropeptide engagement provoke important questions about stability and homeostatic roles for neuromodulation.

Along similar lines, it is also worth considering how selected intracellular signaling cascades (i.e. G-proteins vs arrestins vs kinases) during MOPR-signaling may further contribute to the heterogeneity of MOPR function. Indeed, some reports have indicated that opioid efficacy and bias can tune opposing behavioral phenotypes ^43^. Though likely of critical importance, the ability to isolate these multifaceted features of endogenous MOPR related circuits, ligands, and signaling pathways has been greatly stymied by limitations of technology ^10^. Integrative use of technologies, like classical pharmacology and the single virus CRISPR approaches used in this study, will be useful for expanding our understanding of neuropeptide biology ^10,26,32,44,45^. Continued advances in hardware ^46–48^ and GPCR-based biosensor development ^49,50^ will also be important to provide critical new insights into the spatiotemporal dynamics of endogenous neuropeptide systems and their control over state-dependent behavior.

The LDRN^MOPR^-mNAcSh^Penk^ circuit we isolated here potentiates multiple voluntary consumptive behaviors, and contributes to exogenous opioid reward, opening many new avenues for research on substance abuse and motivated neural circuits. Future work across neuropeptide biology will necessitate expanding circuit, regional, and peptide investigations to include traditional and nontraditional functional roles, as viewing these systems through an evolutionary lens provides insight and alternative perspectives for therapeutic development for neuropsychiatric disorders.

## Methods Animals

Adult (18–35 g) male and female wildtype, preprodynorphin-IRES-Cre (*Pdyn*-Cre), preproenkephalin-IRES-Cre (*Penk*-Cre), *Oprm1* knockout (*Oprm1* KO), *Oprm1* conditional knockout (*Oprm1*^*fl/fl*^), *Oprm1*^*fl/fl*^ x *Penk*-Cre, *Oprm1*^*fl/fl*^ x *Pdyn*-Cre, *Oprm1* KO x *Penk*-Cre, proopiomelanocortin-cre (*POMC*-Cre) and preprodynorphin conditional knockout (*Pdyn*^*fl/fl*)^ mice were group housed, given access to food pellets and water ad libitum, and maintained on a 12 hr:12 hr light:dark cycle (lights on at 7:00 AM). All mice were kept in a sound-attenuated, isolated holding facility one week prior to surgery, post-surgery, and throughout the duration of the behavioral assays to minimize stress. For cell-type conditional deletion, ablation, chemogenetic, and optogenetic experiments, we used Cre-cage and littermate controls. For *Oprm1* KO, *Oprm1*^fl/fl^ and associated crosses, experimental mice were compared to age-matched wild-type or Cre-littermate controls. Unless otherwise noted, animals had *ad libitum* access to food and water. Any variation from these approaches was due to behavioral attrition from off-target injections/implants or headcap failures. The mice were bred at Washington University in Saint Louis or the University of Washington. Where needed, *Oprm1*^fl/fl^ mice were crossed to Ai14-tdTomato mice on C57BL/6 background, bred, and backcrossed for three generations. All animals were drug and test naive, individually assigned to specific experiments as described, and not involved with other experimental procedures. Statistical comparisons did not detect any significant differences between male and female mice, and were therefore combined to complete final group sizes. All animals were monitored for health status daily and before experimentation for the entirety of the study. All procedures were approved by the Animal Care and Use Committee of Washington University, Animal Care and Use Committee of the University of Washington, and conformed to US National Institutes of Health guidelines.

### Tissue processing

Unless otherwise stated, animals were transcardially perfused with 0.1 M phosphate-buffered saline (PBS) and then 40 mL 4% paraformaldehyde (PFA). Brains were dissected and post-fixed in 4% PFA overnight and then transferred to 30% sucrose solution for cryoprotection. Brains were sectioned at 30 mM on a microtome and stored in a 0.01M phosphate buffer at 4°C prior to immunohistochemistry and tracing experiments. For behavioral cohorts, viral expression and optical fiber placements were confirmed before inclusion in the presented datasets.

### RNAscope Fluorescent In Situ Hybridization

Following rapid decapitation of WT, *Oprm1* KO, *Oprm1 KO x Penk-*Cre, *Oprm1*^fl/fl^, *Oprm1*^fl/fl^ x *Penk*-Cre, *Oprm1*^fl/fl^ x *Pdyn*-Cre, or *Penk*-Cre mice brains were rapidly frozen in 100mL −50°C isopentane and stored at −80°C. Coronal sections corresponding to the site of interest or injection plane used in the behavioral experiments were cut at 20uM at −20°C and thaw-mounted onto SuperFrost Plus slides (Fisher). Slides were stored at −80°C until further processing. Fluorescent in situ hybridization was performed according to the RNAscope 2.0 Fluorescent Multiple Kit User Manual for Fresh Frozen Tissue (Advanced Cell Diagnostics, Inc.) as described by Wang et al. (2012). Briefly, sections were fixed in 4% PFA, dehydrated, and treated with pretreatment 4 protease solution. Sections were then incubated for target probes for mouse mu opioid receptor (*Oprm1*, accession number NM_001039652.1, probe region 1135 - 2162), proenkephalin (*Penk*, accession number <u>NM_001002927.2</u>, probe region 106 – 1332), prodynorphin (*Pdyn*, accession number <u>NM_018863.3</u>, probe region 33 - 700), tryptophan hydroxylase 2 (*tph2*, <u>NM_173391.3</u>, probe region 1640 - 2622), vesicular glutamate transporter type 2 (*slc17a6*, <u>NM_080853.3</u>, probe region 86 - 2998), vesicular GABA transporter (slc32a1, <u>NM_009508.2</u>, probe region 894 - 2037), or Cre (<u>KC845567.1</u>, probe region 1058 – 2032) All target probes except mu consisted of 20 ZZ oligonucleotides and were obtained from Advanced Cell Diagnostics. Following probe hybridization, sections underwent a series of probe signal amplification steps followed by incubation of fluorescently labeled robes designed to target the specific channel associated with the probes. Slides were counterstained with DAPI, and coverslips were mounted with Vectashield Hard Set mounting medium (Vector Laboratories. Images were obtained on an Olympus Fluoview 3000 confocal microscope and analyzed with HALO software. To analyze the images, each image was opened in the HALO software. DAPI positive cells were then registered and used as markers for individual cells. An observer blind to the brain tissue origin and probes used then counted the total number of probe labeled cells for each channel separately. A positive cell consisted of an area within the radius of a DAPI nuclear staining that measured at least 3 positive pixels for receptor probes, or 10 total positive pixels for neurotransmitter probes. Two - three separate slices from the NAc or DRN were used for each animal and that total is presented in the data.

### Stereotaxic Surgery

After mice were acclimated to the holding facility for at least seven days, the mice were anaesthetized in an induction chamber (1%-4% isoflurane) and placed into a stereotaxic frame (Kopf Instruments, model 1900) where they were mainlined at 1%-2% isoflurane. For mice receiving viral injections, a blunt needle (86200, Hamilton Company) syringe was used to deliver the vector at a rate of 100 μL/min. The type of virus, injection volume, and stereotaxic coordinates for each experiment are listed in Table S1 interest (NAc medial shell: AP +1.4, ML ± 0.6, DV −4.3; Dorsal raphe nucleus: AP −4.5, ML ± 0.0, DV −2.6). For mice receiving intracranial implants (i.e., cannulas, fiber photometry or optogenetic optic fibers, or GRIN lens), a small hole was drilled above the site of interest (NAc medial shell: AP +1.4, ML ± 0.6, DV −4.3; Dorsal raphe nucleus: AP −4.5, ML ± 0.0, DV −2.6) and the implant was slowly lowered to the coordinates. Cannulas were secured to the skull using one bone screw and super glue (Lang Dental). All other implants were secured using MetaBond (C & B Metabond).

### Food Intake Test (FI)

Behaviorally tested mice were habituated to a 26×26cm chamber for three days for 1hr. Mice were given free access to ∼0.8g of sucrose pellets during this habituation period. After the final habituation day, mice were either left ad libitum, or food deprived (animals still had access to water). Twenty-four hours later, animals were placed into the food intake chamber for 1hr and had free access to sucrose pellets. Total weight of the sucrose pellets was recorded before and after the test session. After testing, food deprived animals were returned to an ad libitum diet. 48 hours later, animals received the other testing condition (order counter-balanced across cages). Each test session was recorded and behavior scored offline using Ethovision. Animals that did not consume more than 0.01g during the food deprived condition were removed from analysis. Animals that did not lose at least 5% bodyweight after food deprivation were also removed from analysis (animals lost ∼11% bodyweight on average).

### Regulated Food Intake Test

Mice were habituated to a Med-Associates operant conditioning box for three days, during which non-contingent deliveries of two sucrose pellets occurred on a VI of 120sec (±30sec) for 30 minutes. On the final habituation day, mice received an i.p. injection of saline to habituate them to the injection process. Mice were then placed back in the home cage for 15 minutes. At the end of 15 minutes, mice were placed into the test apparatus and behavioral testing proceeded. The same procedure was conducted on each test day, for injections of either saline or naloxone (2mg/kg i.p.). Test days were recorded scored offline by an observer blind to the conditions. Drug order and physiological state were counterbalanced across test days and cages for all experiments. Mice that were used in the free intake photometry experiment were also used for the regulated intake experiments.

### Regulated Lickometer Test

Mice were habituated to a Med-Associates operant conditioning box for three days, during which non-contingent presentation of a lickometer, filled with 10% sucrose solution, occurred on a VI of 120sec (±30sec) for 30 minutes. On the final habituation day, mice received an i.p. injection of saline to habituate them to the injection process. Mice were then placed back in the home cage for 15 minutes. At the end of 15 minutes, mice were placed into the test apparatus and behavioral testing proceeded. The same procedure was conducted on each test day, for injections of either saline or naloxone (2mg/kg i.p.). For opto-MOR and ChR2 experiments, mice received 20Hz stimulation (473 nm, 10 ms pulse width, ∼1-3mW light power) upon the first lick of the lickometer and was continued until the lickometer was retracted. Drug order, laser stimulation, and physiological state were all counterbalanced across test days and cages for all experiments.

### Restraint Stress

For Elevated Zero Maze testing (described below), mice were habituated to the testing room for 1 day prior to testing for 1 hour. On the test day, mice were brought to the test room and allowed 30 additional minutes to habituate. After the second habituation, mice were either placed into a 50mL conical tube for 30 minutes (restrained) or left in the home cage (unrestrained). After the restraint period, mice were released from the tube back into the home cage for 30 minutes. Finally mice were tested on the elevated zero maze or food intake experiments, after which they were returned to the home cage.

### Elevated Zero Maze (EZM)

The EZM (Harvard Apparatus, Holliston, MA) was made of grey plastic, 200 cm in circumference, comprised of four 50 cm sections (two opened and two closed). The maze was elevated 50 cm above the floor and had a path width of 4 cm with a 0.5 cm lip on each open section. Room lighting was maintained at 4 lux. Mice were positioned head first into a closed arm, and allowed to roam freely for 7 min. For the naloxone experiment, mice were injected prior to the 30 minute restraint. Mean open arm time was the primary measure of anxiety-like behavior.

### Drug Microinjections and DREADD Experiments

For drug microinjection experiments, animals were habituated for food intake (described below) one week after cannula implantation. DREADD tested mice were allowed to recover for 4 weeks after surgery before habituation. The day before testing, microinjection animals received an infusion of vehicle (ACSF) to habituate them to the microinjection process. Mice were scruffed and a double 32G microinjector was inserted through the guide cannulas. Animals were then placed into a box in which they could freely walk around during the infusion of vehicle. Mice received a total injection of 200uL per/side over the course of 1 minute. Microinjections were then left in the cannulas for an additional minute to ensure complete diffusion of vehicle. Mice were then placed back in the home cage for 15 minutes. At the end of 15 minutes, mice were placed into the test apparatus and behavioral testing proceeded. The same procedure was conducted on each test day, for injections of either vehicle or the selective mu opioid receptor antagonist CTAP (1ug/1uL). On the final day of habituation, DREADD mice received an i.p. injection of saline to habituate them to the injection process. Mice were then placed back in the home cage for 15 minutes. At the end of 15 minutes, mice were placed into the test apparatus and behavioral testing proceeded. The same procedure was conducted on each test day, for injections of either saline or clozapine-N-oxide (CNO, 3mg/kg i.p.) Drug order and physiological state was counterbalanced across test days and cages all experiments.

### Patch-Clamp Electrophysiology

Mice (n = 5; 4-6 months; 2 males) were anesthetized with pentobarbital (50 mg/kg) before transcardial perfusion with ice-cold sucrose cutting solution containing the following (in mM): 75 sucrose, 87 NaCl, 1.25 NaH_2_P0_4_, 7 MgCl_2_, 0.5 CaCl_2_, 25 NaHCO_3_, 306-308 mOsm. Brains were then rapidly removed, and coronal sections 300 μm thick were taken using a vibratome (Leica, VT 1200). Sections were then incubated in aCSF (32°C) containing the following (in mM): 126 NaCl, 2.5 KCl, 1.2 NaH_2_P0_4_, 1.2 MgCl_2_, 2.4 CaCl_2_, 26 NaHCO_3_, 15 glucose, 305 mOsm.. After an hour of recovery, slices were constantly perfused with aCSF (32°C) and visualized using differential interference contrast through a 40x water-immersion objective mounted on an upright microscope (Olympus BX51WI). Whole-cell patch-clamp recordings were obtained using borosilicate pipettes (4–5.5 Ω) back-filled with internal solution containing the following (in mM): 117 Cs-Methanesulfonate, 20 HEPES, 0.4 EGTA, 2.8 NaCl, 5 TEA, 5 ATP, and 0.5 GTP (pH 7.35, 285 mOsm). To assess connectivity between DRN^*Penk*^ and NACsh, voltage clamp recordings were performed from cells located near eYFP-expressing axons within the NAC. 5 ms blue light pulses were delivered through the objective while holding each cell at - 70 mV and +10 mV to assess glutamatergic and GABAergic input, respectively. During voltage clamp recordings, TTX (1 µM) and 4-AP (1 mM) (Sigma) were applied to the bath, and then the AMPA/kainate receptor antagonist, DNQX (10 µM), or the GABA_A_ antagonist, gabazine (10 µM), were applied to test for glutamate and GABA mediated currents, respectively. Data acquisition occurred at 10 kHz sampling rate through a MultiClamp 700B amplifier connected to a Digidata 1440A digitizer (Molecular Devices). Data were processed using Clampfit v11.0.3.03 (Molecular Devices) and analyzed using GraphPad Prism v8.3.0. All tests were two-sided and corrected for multiple comparisons or unequal variance where appropriate.

### In Vivo Fiber Photometry

Fiber photometry recordings were made throughout the entirety of 60-minute Food Intake, 30-minute Regulated Food Intake, and 30-minute Lickometer Tests. Prior to recording, an optic fiber was attached to the implanted fiber using a ferrule sleeve (Doric, ZR_2.5). Two LEDs were used to excite GCaMP6s. A 531-Hz sinusoidal LED light (Thorlabs, LED light: M470F3; LED driver: DC4104) was bandpass filtered (470 ± 20 nm, Doric, FMC4) to excite GCaMP6s and evoke Ca^2+^-dependent emission. A 211-Hz sinusoidal LED light (Thorlabs, LED light: M405FP1; LED driver: DC4104) was bandpass filtered (405 ± 10 nm, Doric, FMC4) to excite GCaMP6s and evoke Ca^2+^-independent isosbestic control emission. Prior to recording, a 120 s period of GCaMP6s excitation with 405 nm and 470 nm light was used to remove the majority of baseline drift. Laser intensity for the 470 nm and 405 nm wavelength bands were measured at the tip of the optic fiber and adjusted to ∼50 μW before each day of recording. GCaMP6s fluorescence traveled through the same optic fiber before being bandpass filtered (525 ± 25 nm, Doric, FMC4), transduced by a femtowatt silicon photoreceiver (Newport, 2151) and recorded by a real-time processor (TDT, RZ5P). The envelopes of the 531-Hz and 211-Hz signals were extracted in real-time by the TDT program Synapse at a sampling rate of 1017.25 Hz.

### Photometry Analysis

Custom MATLAB scripts were developed for analyzing fiber photometry data in context of mouse behavior and can be accessed via GitHub. The isosbestic 405 nm excitation control signal was subtracted from the 470 nm excitation signal to remove movement artifacts from intracellular Ca^2+^-dependent GCaMP6s fluorescence (see Figure S3B). Baseline drift was evident in the signal due to slow photobleaching artifacts, particularly during the first several minutes of each hour-long recording session. A double exponential curve was fit to the raw trace and subtracted to correct for baseline drift. After baseline correction, the photometry trace was z-scored relative to the mean and standard deviation of the test session. The post-processed fiber photometry signal was analyzed in the context of animal behavior during food intake and lickometer tests.

### Tail-immersion Test

Tail-immersion tests were performed by submerging 2-4cm of the mouse tail into hot water (54°C) and recording the total time between tail immersion and withdrawal. The cut-off was 10sec after immersion in order to prevent permanent damage to the tail tissue. After the initial immersion response (time 0), mice were injected with morphine (5mg/kg, s.c.) and tested every 10 minutes for 1 hour, then again at 90 minutes.

### Morphine Conditioned Place Preference (CPP)

Mice were trained in an unbiased, balanced three compartment conditioning apparatus as previously described (McCall et al., 2015). Briefly, mice were pre-tested by placing individual animals in the small central compartment and allowing them to explore the entire apparatus for 30 min. Time spent in each compartment was recorded with a video camera (ZR90; Canon, Tokyo, Japan) and analyzed using Ethovision 11 (Noldus). The drug paired side was randomly assigned to the mice. For the three conditioning days, mice received a subcutaneous injection of saline in the morning, and six hours later received an injection of morphine (5mg/kg, s.c.). CPP was assessed on day 5 by allowing the mice to roam freely in all three compartments and recording the time spent in each. The difference in preference for context A and B from the posttest to the pretest was used as the ultimate preference score.

### In vivo Ca^2+^ Imaging Behavior

Mice were habituated to the food intake chamber for 3 days. During these habituation sessions, they were also mounted with an Inscopix miniature microscope (nVista) that was attached to a commutator to prevent the cord from tangling. Prior to the first test day, scope focus, power, and gain were optimized individually for each mouse. These settings were then kept constant for each test day. During test days, mice were allowed to freely consume sucrose pellets for 30 minutes, during which GCaMP6s fluorescence was recorded. When the test day was over, mice were returned to the home cage.

### In vivo Ca^2+^ Imaging Data Processing

Inscopix data acquisition software (IDAS; Inscopix) was used to acquire TIFF images of fluorescence dynamics at 20 frames per second. Calcium frame acquisition was triggered via Ethovision XT (v10) and began at the onset of the session. Calcium acquisition was automatically terminated after 30 minutes.

Inscopix data processing software (IDPS; Inscopix) was used to preprocess Ca^2+^ data from each imaging session as previously described (*102, 103*). Briefly, for each day, calcium imaging data from *ad libitum* and food deprived test days were down-sampled temporally (2x temporal bin) and spatially (4x spatial bin) and rigid motion correction was applied. After preprocessing, putative single neuron activity was segmented using Constrained Non-negative Matrix Factorization for Endoscopic data (CNMFe) using custom MATLAB (MathWorks, Natick, MA, USA) scripts. Individual putative striatal neurons were tracked between *ad libitum* and food deprived sessions using CellReg (cite PMID: 29069591). For each mouse the spatial correlation registration threshold performed best, and thresholds were determined by the algorithm. Final registration utilized the probabilistic model with a P_same (probability of cells being the same) > 0.6 for all mice.

### *In Vivo* Ca^2+^ Imaging Data Analysis

Raw fluorescence traces for individual putative neurons were Z-normalized 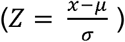 using the mean and standard deviation of the entire imaging session. Individual neurons were grouped together based on similar activity during approach and consumption of sucrose rewards using a standard k-means clustering approach. The optimal number of clusters was determined by silhouette analysis. Discrete timestamps for the onset of sucrose consumption, sniffing and rearing were manually scored from simultaneously recorded behavioral videos. The mean activity of neuronal clusters was then plotted relative to the onset of behavioral events. Normalization and analysis of individual neuron calcium dynamics was performed in MatLab using custom scripts.

### Generation and validation of AAV1-FLEX-SaCas9-U6-sg*Penk*

The sgRNA targeted to the *Penk* locus (sg*Penk*) was designed as previously described (Hunker et al., 2020). The following oligos (Sigma) were used to clone into pAAV-FLEX-SaCas9-U6-sgRNA (Addgene Cat#: 124844). *Penk* forward: CACCGTTTGCACCTGGCTGCTGGCGC; *Penk* reverse: AAACGCGCCAGCAGCCAGGTGCAAAC.

### Targeted deep sequencing of *Penk* locus

Nuclei isolation, FACS, and targeted deep sequencing were performed as described previously (Hunker et al., 2020). Tissue punches of the ventral striatum from 4 mice co-injected with AAV1-FLEX-SaCas9-U6-sg*Penk* and AAV1-FLEX-EGFP-KASH were pooled into a single group and homogenized in 2mL of homogenization buffer containing (in mM): 320 Sucrose (sterile filtered), 5 CaCl (sterile filtered), 3 Mg(Ac)2 (sterile filtered), 10 Tris pH 7.8 (sterile filtered), 0.1 EDTA pH 8 (sterile filtered), 0.1% NP40, 0.1 Protease Inhibitor Cocktail (PIC, Sigma Cat#: P8340), 1 β-mercaptoethanol. Homogenization was performed using 2mL glass dounces (Sigma Cat#: D8938-1SET); 25 times with pestle A, then 25 times with pestle B. The volume of the homogenate was transferred to a 15mL conical tube and brought up to 5mL using homogenization buffer, mixed by inversion, and incubated on ice for 5 minutes. 5mL of 50% Optiprep density gradient medium (Sigma Cat#: D1556-250ML) containing (in mM): 5 CaCl (sterile filtered), 3 Mg(Ac)2 (sterile filtered), 10 Tris pH 7.8 (sterile filtered), 0.1 PIC, 1 β-mercaptoethanol was added to the homogenate and mixed by inversion. The mixture was gently loaded on 10mL of 29% iso-osmolar Optiprep solution in a 1×3 1/2 in Beckman centrifuge tube (SW32 Ti rotor) and spun at 7500 RPM for 30min at 4°C. The floating cell debris was removed using a KimWipe and the supernatant was gently poured out. The nuclei pellet was vigorously resuspended in sterile 1xPBS. 500 GFP-positive and 500 GFP-negative nuclei were sorted directly into 3uL of REPLI-g Advanced Storage buffer (Qiagen Cat#: 150365) in an PCR tube strip (Genessee Cat #: 24-706) using a BD AriaFACS III. Whole genome amplification (WGA) was performed directly following FACS using the REPLI-g Advanced DNA Single Cell kit (Qiagen Cat#: 150365) according to manufacturer’ s instructions.

For generation of the specific amplicons, 1ul of WGA DNA was diluted 1:50 and amplified (PCR 1) with Phusion High Fidelity Polymerase (Thermo Fisher Cat#: F530L) using the following thermocycler protocol: initial denaturation (30sec, 95°C); denaturation (10sec, 95°C); annealing (20sec, 66°C); extension (10sec, 72°C); cycle repeated x34; final extension (5min, 72°C). PCR forward: GCTCAGGAAAGACTGTCC, PCR reverse: TGACCACTAGAAGTCTGC. 1uL of PCR 1 was amplified again (PCR 2) with the same set of primers using the same thermocycler protocol. The 310bp amplicon from PCR 2 was gel extracted using the MinElute gel extraction kit (Qiagen Cat#: 28606) and sent to Genewiz for Amplicon-EZ targeted deep sequencing and Sanger sequencing. Reads received from Amplicon-EZ were trimmed in Excel up to the sgRNA and PAM sequence to avoid false mutational reads due to PCR error. The number of unique reads containing specific insertions, deletions, and base changes within the targeted region were then summed in Excel.

### Statistical analyses

All data collected were averaged and expressed as mean ± SEM. Statistical significance was taken as ∗p < 0.05, ∗∗p < 0.01, and ∗∗∗p < 0.001, as determined by Pearson’ s correlation, Student’ s t test, one-way ANOVA or a two-way repeated-measures ANOVA followed by Tukey post hoc tests as appropriate. For *in situ* hybridization and electrophysiology data, we used Student’ s t test. For photometry experiments, we used Pearson’ s correlation and Student’ s t tests, as appropriate. For microinjection, genetic deletion, photometry, 1-photon, ablation, chemogenetic, and optogenetic behavioral experiments, we used one-way or two-way repeated-measures ANOVA followed by a Tukey post hoc tests. All n values for each experimental group are described in the appropriate figure legend. For behavioral experiments, group size ranged from n = 3 to n = 15. For *in situ* hybridization quantification experiments, slices were collected from 2-3 mice, with data averaged from 2-3 slices per mouse. For electrophysiology experiments, the number of cells recorded were as follows: n = 107 total recorded cells, 8 cells that showed oEPSPs, and 5 cells that showed oIPSPs. Statistical analyses were performed in GraphPad Prism 8.0 (Graphpad, La Jolla, CA) and MATLAB 9.6 (The MathWorks, Natick, MA).

## Data and Code Availability

Custom MATLAB analysis and code was created to appropriately organize, process, and combine photometry and single-photon recording data with associated behavioral data. Analysis code for photometry and single-photon imaging from Figures 3 and 5 will be made available on Github. The behavioral dataset supporting the current study are available from the corresponding author upon request.

## Acknowledgements

We thank Lamley Lawson, Dylan Blumenthal, Michelle Chung, Taylor Hobbs, and Carina Pizzano for animal colony maintenance. We thank the Bruchas lab, Stuber lab and NAPE Center for helpful discussions.

## Author Contributions

D.C.C designed and performed experiments, collected and analyzed data, and wrote the manuscript. C.O., E.T.Z., and A.G. performed experiments and collected data. M.A.R. collected and analyzed electrophysiological data. A.H. and L.S.Z. designed and analyzed CRISPR virus. C.E.P designed and analyzed fiber photometry data. S.C.P. designed and analyzed 1-photon data. M.R.B., and J.M.C facilitated resources for generation of *Oprm1*^*fl/fl*^ x *Penk*/*Pdyn* mouse lines. J.M.C, L.S.Z., and G.D.S helped to design experiments, discuss results and write the manuscript. M.R.B helped lead the design, analysis, oversight of experiments, discuss results, provide resources, and write the manuscript.

## Data and materials availability

Plasmids generated in this study will be deposited to Addgene. Mouse lines generated in this study will be deposited to Jackson Laboratories. This study did not generate any new/unique reagents. Custom BATLAB analysis and code was created to appropriately organize, process, and combine photometry and single-photon recording data with associated behavioral data. Analysis code for photometry and single-photon imaging from Figures 3 and 5 will be made available on Github. The behavioral dataset supporting the current study are available from the corresponding author upon request.

## Materials and Correspondence

Further information and requests for resources and reagents should be directed to and will be fulfilled by corresponding author Michael R. Bruchas.

