## Extended Data for "An endogenous opioid circuit determines state-dependent appetitive behavior"

**Title: An endogenous mu-opioid circuit determines**

**state-dependent appetitive behavior**

**This document includes**

Extended Data Text

Extended Data Figures 1-10

Extended Data Tables 1-2

**Other Supplementary Materials for this manuscript include the following:**

Movie S1

**Extended Data Text**

***Endogenous MOPRs mediate avoidance motivation, but via dissociable mechanisms***

Alongside the state-dependent sucrose consumption task, we tested mice in an oppositional avoidance task, the elevated zero-maze (EZM, **Fig. S2A-C**), to evaluate whether MOPRs can potentiate both avoidance-type and appetitive-type behaviors. We found that in wildtype mice, acute restraint stress significantly potentiated avoidance behavior for the open arms of the maze, as previously described (*37*, *38*), and that systemic pre-treatment with the opioid antagonist naloxone (2mg/kg, i.p.) or MOPR deletion in *Oprm1* KO mice prevented restraint enhanced avoidance. This result suggests that MOPRs are involved in potentiating both appetitive and avoidance behaviors in response to acute psychophysiological stress (i.e., food deprivation or restraint). However, *OPRM1^fl/fl^* x *Penk*-Cre^+^ mice still had significant avoidance after restraint stress, suggesting that while MOPRs may broadly be involved in both appetitive and avoidance behaviors, these behaviors are relegated by separable neuronal populations. Further support for this conclusion can be drawn by an experiment whereby we restrained mice (i.e., recruited endogenous opioids) and tested them on the sucrose consumption task (*30*, *39*) (**Fig. S2D-E**). We found that restraint was unable to potentiate food intake, indicating that specific MOPR regulation of appetitive behaviors is controlled by discrete neurophysiological mechanisms.

***Additional in situ analysis of the LDRN^Penk^-mNAcSh projection***

FISH experiments revealed that MOPRs were more likely to coexpress with vGAT neurons (27% total, 12% overlap) over lateralized vGlut2 populations (14% total, 6% overlap), suggesting that LDRN^Penk^ populations, which are also lateralized, are likely GABAergic (**Fig. S3A-C**). To test whether DRNPenk-mNAcSh cells were additionally MOPR expressing, we injected a retrograde virus into mNAcSh to label incoming neurons with Cre. FISH analyses in DRN revealed that ~60% of cre-labeled (i.e., mNAcSh projecting) enkephalin neurons also expressed MOPR (**Fig. S3D-G**). Similarly, ~60% of cre-labeled MOPR expressing cells also expressed enkephalin. As a non-viral retrograde tracing method control (to exclude differences due to tropism), we also injected fluorescently tagged CTb into mNAcSh (**Fig. S3H-I**). Consistent with our retrograde viral approach, we found CTb labeled cells in LDRN, further establishing that LDRN projects to mNAcSh.

***Non-LDRN^Penk^-mNAcSh populations are not necessary for potentiated sucrose consumption***

Several regions, such as basomedial amygdala (BMA), paraventricular thalamus (PVT), and ventral pallidum (VP) were also found to fluorescently labeled (**Extended Data Fig. 3A). To test whether these regions were functionally necessary for food deprived potentiation** of sucrose consumption, we first deleted MOPRs in PVT neurons in *Oprm1^fl/fl^* mice. Prior work has shown that PVT-NAc projections modulate appetitive behavior, and PVT, like mNAcSh, is enriched in MOPR. However, we did not observe any significant disruptions in food deprived sucrose intake after PVT MOPR deletion (Extended Data Fig. 4A). We also ablated enkephalin neurons, and by extension potential MOPR-expressing populations, in BMA or VP using a caspase approach (Extended Data Fig. 4B-D). Similar to the PVT^MOPR^ deletion experiment, caspase ablation of neither BMA^Penk^ nor VP^Penk^ disrupted food deprived sucrose intake as compared to Penk-Cre^-^ controls.

***MOPR activation on LDRN^Penk^ does not restore morphine analgesia***

To further test the generalizability of LDRN^Penk^ in regulating opioid-dependent behavior, we examined an additional behavior where MOPR is necessary, morphine analgesia. Previous studies have shown that *Oprm1* KO mice do not show morphine analgesia, as measured by a tail-immersion test. Here, the tails of mice were dipped into hot water (52ºC) and the latency to flick the tail out of the water (nociceptive response) was recorded before and after morphine administration (5mg/kg, s.c.). Consistent with many previous reports, wildtype mice showed a short latency (~3sec) to flick their tail out of the water prior to morphine administration (Extended Data Fig. 6B-C). Following morphine, the tail flick latency was significantly increased to the maximum allowable immersion time (10sec). In contrast, *Oprm1* KO x *Penk*-Cre^-^ mice showed similar tail-flick latencies as compared to wildtype controls prior to morphine administration but did not show an analgesic response after morphine. Finally, *Oprm1* KO x *Penk*-Cre^+^ with viral MOPR rescue in LDRN^Penk^ also showed similar baseline tail-flick latencies to wildtypes, but like *Penk*-Cre^-^ mice, did not show morphine analgesia (Extended Data Fig. 6C). These data indicate that despite its proximity to the pain-associated periaqueductal gray, MOPRs in LDRN^Penk^ do not act to regulate morphine analgesia.

***Physiological Inhibition of LDRN^MOPR^-mNAcSh is necessary for potentiation of appetitive behavior***

We next sought to examine whether MOPR-mediated inhibition of LDRN^MOPR^-mNAcSh terminals was required to potentiate sucrose consumption behaviors. To test this type of physiological necessity, disrupted MOPR inhibition by stimulating the ChR2 expressing LDRN^Penk^ neurons, terminals in mNAcSh during the sucrose solution task (Extended Data Fig. 6E). We found that ChR2 stimulation in the *ad libitum* state did not disrupt baseline intake. (Extended Data Fig. 6F). In contrast, photo-stimulation of LDRN^MOPR^-mNAcSh terminals in the water deprived state significantly reduced overall licks by ~50% and ChR2 stimulation appeared to disrupt the engagement of long, sustained licking bouts (Extended Data Fig. 6G-H). These data indicate that temporally precise engagement of opioid inhibition in the LDRN^MOPR^-mNAcSh circuit is necessary for the potentiation of appetitive behaviors.

***Arcuate POMC neurons do not modulate potentiated sucrose consumption***

To test for other sources of endogenous MOPRs agonists, and to determine if there is a unique role local mNAcSh^Penk^ in activating MOPRs on LDRN^Penk^-mNAcSh terminals, we also expressed HM3D(Gq) or HM3D(Gi) DREADDs into the arcuate nucleus of *POMC*-Cre^+^ or *POMC*-Cre^-^ mice (Extended Data Fig. 8C-G). *POMC* is the precursor peptide for the most selective MOPR ligand, beta-endorphin, and is only produced within the arcuate nucleus of the hypothalamus or nucleus of the solitary tract in the brainstem. We found that neither Gq-DREADD nor Gi-DREADD stimulation of arcuate *POMC*-Cre^+^ neurons modulated sucrose consumption in *ad libitum* or food deprived states, suggesting that beta-endorphin producing neurons are not a likely endogenous source for MOPR agonism in mNAcSh for opioid-mediated potentiation of sucrose consumption.

***Local mNAcSh^Pdyn^ does not modulate potentiated sucrose consumption***

Consistent with our initial FISH analyses of mNAcSh, we found that there was a sizeable overlap between *Pdyn* and *Penk* labeled mRNA in this subregion (Fig. 1H, 6C). It is therefore unsurprising that in our analysis of the caspase treated mice, we observed reductions in both *Pdyn* and *Penk* expression, since at least some of the *Pdyn^+^* cells were also *Penk*-Cre^+^ (Fig. 6C). It is possible, then, that the effects we observed in the DREADD and caspase experiments could be driven by local mNAcSh^Pdyn^ and/or the MOPR agonist Leu-enkephalin (i.e., Dyn 1-8) release, rather than by met-enkephalin (produced by *Penk* neurons) (*59*). To test this possibility, we injected AAV5-CMV-Cre-GFP into mNAcSh of *Pdyn^fl/fl^* mice to delete dynorphin production in this region and tested mice in the voluntary sucrose consumption paradigm (Extended Data Fig. 9C). Unlike DREADD and caspase treated mice, *Pdyn^fl/fl^* mice did not reduce food deprived sucrose intake, indicating that local mNAcSh dynorphin and/or leu-enkephalin production is not necessary for LDRN^MOPR^-mNAcSh associated appetitive behavior.
