## Extended Data Table 1 for "An endogenous opioid circuit determines state-dependent appetitive behavior"

**Extended Data Table 1. Key Resources**

| REAGENT or RESOURCE | SOURCE | IDENTIFIER |
| --- | --- | --- |
| Antibodies | | |
| Chicken-anti-GFP | Abcam | Ab13970 |
| Anti-Chicken 488 | Abcam | Ab150169 |
| Bacterial and Virus Strains | | |
| rAAV5-CMV-Cre-GFP | The Hope Center Viral Core – Washington University at St. Louis | N/A |
| AAV2retro-CMV-myc-NLS-Cre | The Hope Center Viral Core – Washington University at St. Louis | N/A |
| AAV2retro-GFP-Cre | The Hope Center Viral Core – Washington University at St. Louis | N/A |
| AAV2retro-EF1α-DIO-EYFP | The Hope Center Viral Core – Washington University at St. Louis | N/A |
| AAV5-EF1α-ChR2-EYFP | The Hope Center Viral Core – Washington University at St. Louis | N/A |
| AAV5-DJ-EF1α-DIO-GCAMP6s | Stanford University Gene Vector and Viral Core | N/A |
| AAV5-EF1α-DIO-rMOR-GFP | The Hope Center Viral Core – Washington University at St. Louis | N/A |
| AAV5-EF1α-DIO-oMOR-EYFP | The Hope Center Viral Core – Washington University at St. Louis | N/A |
| AAV5-hsyn-DIO-HM4D(Gq)-mCherry | Addgene | N/A |
| AAV5-hsyn-DIO-HM3D(Gi)-mCherry | addgene | N/A |
| AAV2-FLEX-taCas3-TEVp | UNC Vector Core | N/A |
| AAV1-CMB-FLEX-Sa-Cas9U6-sg*Penk* | This Manuscript | N/A |
| Chemicals, Peptides, and Recombinant Proteins | | |
| CTAP | Tocris | CAT#1560; CAS: 103429-32-9 |
| VECTASHIELD Hardset Antifade Mounting Medium | Vector Laboratories | CAT#H-1400 |
| VECTASHIELD Hardset Antifade Mounting Medium with DAPI | Vector Laboratories | CAT#H-1800 |
| Clozapine-N-Oxide | Enzo Sciences | CAT#BML-NS105-0025; CAS: 34233-69-7 |
| Naloxone hydrocholoride | Tocris | CAT#0599; CAS: 357-08-4 |
| Morphine | Tocris | CAT#5158 ; CAS: 52-26-6 |
| Critical Commercial Assays | | |
| RNAscope Fluorescent Multiplex Kit 2.0 | Advanced Cell Diagnostics | CAT#320850 |
| Mm-*OPRM1* | Advanced Cell Diagnostics | CAT#315841 |
| Mm-*Pdyn* | Advanced Cell Diagnostics | CAT#318771 |
| Mm-*Penk* | Advanced Cell Diagnostics | CAT#318761 |
| Mm-TPH2 | Advanced Cell Diagnostics | CAT#318691 |
| Mm-Slc17a6 | Advanced Cell Diagnostics | CAT#319171 |
| Mm-Slc32a1 | Advanced Cell Diagnostics | CAT#319191 |
| Mm-Cre | Advanced Cell Diagnostics | CAT#312281 |
| Experimental Models: Organisms/Strains | | |
| *Oprm1* KO | Jackson Laboratories | Stock No: 007559 |
| *Oprm1^fl/fl^* | (*99*) | Stock No: 030074 |
| *Pdyn*-IRES-Cre | Gift from Dr. Richard Palmiter | Stock No: 027958 |
| *Penk*-IRES-Cre | (*101*) | Stock No: 025112 |
| *Pdyn*-Cre x *Oprm1^fl/fl^* | This paper | N/A |
| *Penk*-Cre x *Oprm1^fl/fl^* | This paper | N/A |
| *Oprm1* KO x *Penk*-Cre | This paper | N/A |
| *POMC*-Cre | Jackson Laboratories | Stock No: 005965 |
| *Pdyn*-Cre^fl/fl^ | Bred in house | Gift from Charley Chavkin |
| Software and Algorithms | | |
| FIJI/ImageJ | NIH | <https://fiji.sc/> |
| MATLAB | Mathworks | <https://www.mathworks.com/products.html> |
| Med-PC V Software Suite | Med-Associates Inc. | <https://www.med-associates.com/med-pc-v/> |
| Ethovision 10 | Noldus | <https://www.noldus.com/ethovision-xt> |
| Synapse | Tucker-Davis Technologies | <https://www.tdt.com/files/manuals/SynapseManual.pdf> |
| PRISM 8 | Graphpad | <https://www.graphpad.com/> |
| Photometry analysis code | Parker et al. | <https://github.com/BruchasLab> |
| Clampfit v11.0.3.03 | Molecular Devices | https://www.moleculardevices.com/products/axon-patch-clamp-system/acquisition-and-analysis-software/pclamp-software-suite |
| Olympus Fluoview 3000 | Leica Microsystems | <https://www.leica-microsystems.com/products/microscope-software/p/leica-las-x-ls/> |
| Illustrator CS6 | Adobe | https://www.adobe.com/products/illustrator.html |
| nVista | Inscopix | https://www.inscopix.com/nVista |
