## Extended Data Table 2 for "An endogenous opioid circuit determines state-dependent appetitive behavior"

**Extended Data Table 2. Overview of Mouse Lines and Experiments**

| Mouse Line | Behavior | Approach | Virus |
| --- | --- | --- | --- |
| *Oprm1* KO | Sucrose Consumption (Figure 1) | N/A | N/A |
|  | Sucrose Consumption (Figure 3) | Viral Injection | AAV5-EF1α-DIO-rMOR-GFP |
|  | Morphine CPP (Figure 3) | Viral Injection | AAV5-EF1α-DIO-rMOR-GFP |
| *Oprm1* KO x *Penk*-Cre | Sucrose Consumption (Figure 3) | Viral Injection | AAV5-EF1α-DIO-rMOR-GFP |
| *Oprm1^fl/fl^* | Sucrose Consumption (Figure 1) | Viral Injection  Viral Injection  Viral Injection | rAAV5-CMV-Cre-GFP  AAV2retro-GFP-Cre  AAV2retro-CMV-myc-NLS-Cre |
| *Pdyn*-Cre x *Oprm1^fl/fl^* | Sucrose Consumption (Figure 1) | Genetic Cross | N/A |
| *Penk*-Cre x *Oprm1^fl/fl^* | Sucrose Consumption (Figure 1) | Genetic Cross | N/A |
| *Penk*-IRES-Cre | Anatomical Tracing (Figure 2) | Viral Injection  Viral Injection  Viral Injection, Patch Clamp | AAV2retro-EF1α-DIO-EYFP AAV2retro-GFP-Cre AAV5-EF1α-ChR2-EYFP |
|  | Sucrose Consumption (Figure 2) | Viral Injection, Fiber Photometry | AAV5-DJ-EF1α-DIO-GCAMP6s |
|  | Sucrose Solution Consumption (Figure 3) | Viral Injection, Optogenetics | AAV5-EF1α-DIO-oMOR-EYFP |
|  | Sucrose Consumption (Figure 4) | Viral Injection, 1-Photon Imaging | AAV5-DJ-EF1α-DIO-GCAMP6s |
|  | Sucrose Consumption (Figure 5) | Viral Injection, DREADD | AAV5-hsyn-DIO-HM4D(Gq)-mCherry  AAV5-hsyn-DIO-HM3D(Gi)-mCherry |
|  | Sucrose Consumption (Figure 5) | Viral Injection, Caspase | AAV2-FLEX-taCas3-TEVp |
|  | Sucrose Consumption (Figure 5) | Viral Injection, CRISPR | AAV1-CMB-FLEX-Sa-Cas9U6-sg*Penk* |
